# Membranous translation platforms in the chloroplast of *Chlamydomonas reinhardtii*

**DOI:** 10.1101/2024.07.22.604633

**Authors:** Yi Sun, Shiva Bakhtiari, Melissa Valente-Paterno, Heng Jiang, William Zerges

**Affiliations:** Department of Biology, Concordia University, 7141 Sherbrooke W, Montreal, Quebec, H4B 1R6, Canada; current address: Department of Biology, University of Oxford, South Parks Road, Oxford, OX1 3RB, United Kingdom; current address: Department of Anatomy and Cell Biology, McGill University, 3640 University, Montreal, Quebec, H3A 0C7, Canada; Centre for Biological Applications of Mass Spectrometry, Concordia University, 7141 Sherbrooke W, Montreal, Quebec, H4B 1R6, Canada

**Keywords:** ribosome, membrane biogenesis, translation, photosystem, chlorophyll, chloroplast, Chlamydomonas

## Abstract

A small genome in chloroplasts encodes polypeptide subunits of the photosynthetic electron transport complexes in the membranes of thylakoid vesicles in the chloroplast stroma. Trans-membrane subunits of these complexes undergo co-translational membrane insertion during their synthesis by ribosomes of the bacterial-like genetic system of this semiautonomous organelle. While thylakoid membranes are sites of translation, evidence in the unicellular alga *Chlamydomonas reinhardtii* supports translation also on non-canonical membranes in a discrete translation-zone in the chloroplast. To characterize the membranous platforms for translation and the biogenesis of thylakoid membrane, we profiled membranes during chloroplast development, using the *yellow-in-the-dark 1* mutant, and carried out proteomic analyses on membranes of interest. The results support roles of two membrane types in preliminary and ongoing stages of translation: a “low-density membrane” and a denser “chloroplast translation membrane”, respectively. These roles are based on correlations of the relative levels of each membrane type and the translational status of the chloroplast before, during and after chloroplast differentiation and results of proteomic analyses. Our results support a model of photosynthesis complex biogenesis in a spatiotemporal “assembly line” involving LDM and CTM as sequential stages leading to photosynthetic thylakoid membranes.

## INTRODUCTION

Chloroplasts have some three thousand proteins for their biogenesis, homeostasis, and intracellular functions, such as photosynthesis. Most chloroplast proteins are encoded by the nuclear genome, synthesized in the cytosol by 80S ribosomes, and then imported across the dual membranes of the chloroplast envelope (Rochaix, 2023). Other chloroplast proteins are encoded by the chloroplast genome and synthesized by the bacterial-type 70S ribosomes of this semi-autonomous organelle (Sun and Zerges, 2015). Among these are integral membrane protein subunits of the photosynthetic electron transport (PET) complexes and ATP synthase, which are embedded in the membranes of the thylakoid vesicles within the chloroplast stroma (Ostermeier et al., 2024). These subunits are integrated into a membrane during their synthesis (i.e. co-translationally). The recipient includes thylakoid membranes (TM), at least for the translation of the *psbA* mRNA in the repair of photodamaged photosystem II (PSII) (Järvi et al., 2015). This is the predominant mode of translation in chloroplasts of mature plant tissues in the light, such as those used in most studies of TM-localized translation (Kettunen et al., 1997; Nilsson et al., 1999; Sun and Zerges, 2015; Walter et al., 2015). The de novo biogenesis of the photosynthesis complexes requires the translation of multiple mRNAs to synthesize multiple subunits and occurs in chloroplasts that are undergoing differentiation or growth and division in developing green tissues (Zoschke and Barkan, 2015; Chotewutmontri and Barkan, 2016). While it is widely believed that such biogenic translation also occurs on TM, recent evidence supports roles for non-canonical chloroplast membranes in *Chlamydomonas reinhardtii*. In the single chloroplast of this unicellular alga, a subcompartment, called the “translation zone” (T-zone), is specialized in the synthesis and assembly of photosystem subunits and chlorophyll and is more discretely localized than the TM throughout the chloroplast (Uniacke and Zerges, 2007; Sun et al., 2019). This suggests the existence of a membranous platform for translation and photosystem biogenesis distinct from TM. Biochemical analyses revealed two types of membrane with chloroplast ribosomes and biogenic factors: “low density membrane” (LDM) and “chloroplast translation membrane” (CTM), which have lower and higher buoyant densities than TM, respectively (Zerges and Rochaix, 1998; Schottkowski et al., 2012). Roles of LDM and CTM in TM biogenesis are based on their cofractionation with a few marker proteins of translation by chloroplast ribosomes and photosystem biogenesis (Zerges and Rochaix, 1998; Schottkowski et al., 2012).

Here, we report comprehensive evidence of LDM and CTM roles in distinct phases of TM biogenesis. This includes changes in LDM and CTM abundances with the translation for TM biogenesis in the chloroplast before, during and after chloroplast differentiation, and the relative abundances of proteins involved in biogenesis versus photosynthesis. Our results highlight how diverse non-canonical chloroplast membranes can be revealed by the analyses of membranes resolved across a broad buoyant density range (Schottkowski et al., 2012). Finally, we provide evidence of spatial coordination between chloroplast and cytoplasmic translation systems in the synthesis and targeting of chloroplast proteins (Billakurthi and Loudya, 2024; Sun et al., 2024).

## RESULTS

### Membranes with chloroplast ribosomes differ with stage of chloroplast development

The abundance of a membranous platform for translation by chloroplast ribosomes should positively correlate with the levels of translation and biogenesis in the chloroplast. To test this prediction, we profiled the relative levels of LDM, CTM and TM before, during and after chloroplast differentiation of *yellow-in-the-dark 1* (*y1*) cells (Ohad et al., 1967; Malnoe et al., 1988). When *y1* is cultured in the dark, on acetate instead of photosynthesis, these “dark-*y1* cells” have an undifferentiated plastid lacking photosystems and TM due to their deficiency of a chlorophyll biosynthetic enzyme, light-independent protochlorophyllide-oxido-reductase. Upon illumination, dark-*y1* cells commence chlorophyll biosynthesis using a light-dependent protochlorophyllide-oxido-reductase (Cahoon and Timko, 2000). Over the subsequent 12 hours in the light, these cells undergo TM differentiation as they carry out enhanced rates of translation for *de novo* photosystem biogenesis (Malnoe et al., 1988; Sun et al., 2019). Hence, dark-*y1* cells are poised for high levels of chloroplast translation. After 2 h of illumination, “2h-*y1* cells” undergo high levels of chloroplast translation for photosystem biogenesis. Ater 16 hours of illumination, the “16h-*y1* cells” are phenotypically wild-type and undergo chloroplast translation for growth and homeostasis (Sun and Zerges, 2015; Sun et al., 2019).

We compared the levels of LDM, CTM and TM before, during and after chloroplast differentiation by resolving membranes from dark-*y1*, 2h-*y1* and 16h-*y1* cells by buoyant density, and then testing the fractions for relevant marker proteins (Figure 1). Post-greening, 16h-*y1* cells yielded abundant TM, seen as a band with the highest levels of PSI and PSII and chlorophyll, the major photopigment of the photosystems and their light-harvesting complexes (LHC) (Figure 1 A and D, F4) (Ohad et al., 1967). The 16h-*y1* cells also yielded CTM in F5-F7, as detected by the presence of markers for the 30S and 50S subunits of the chloroplast ribosome, low PSI and PSII levels, and higher buoyant density relative to TM (Figure 1 D) (Schottkowski et al., 2012). Dark- *y1* cells yielded an abundant yellow membrane as a discrete band with higher density than TM (Figure 1 A, B, and D) (Ohad et al., 1967). This band was CTM because it had the highest levels of 30S and 50S chloroplast ribosomal subunits and was denser than the TM in the gradient of 16h-*y1* cell membranes (Figure 1 B and D) (Schottkowski et al., 2012). This band also had the trace amounts of the PSI and PSII subunits that accumulate under these conditions, consistent with the proposed role of CTM in TM protein synthesis (Schottkowski et al., 2012). Chloroplast ribosomes on CTM were likely translating because dark-*y1* cells synthesize photosystem subunits, albeit at lower rates compared to greening 2h-*y1* cells (Klein et al., 1988). High rates of chloroplast translation in greening 2h-*y1* cells correlated with a prominence of CTM; membranes across all densities had both chloroplast ribosome subunits and RBP40, a protein required for translation of the chloroplast *psbD* mRNA (Figure 1 C) (Schwarz et al., 2007). These results reveal that the activation of chloroplast translation during *y1* greening is accompanied by increases in CTM abundance and range of buoyant densities.

**Figure 1.**
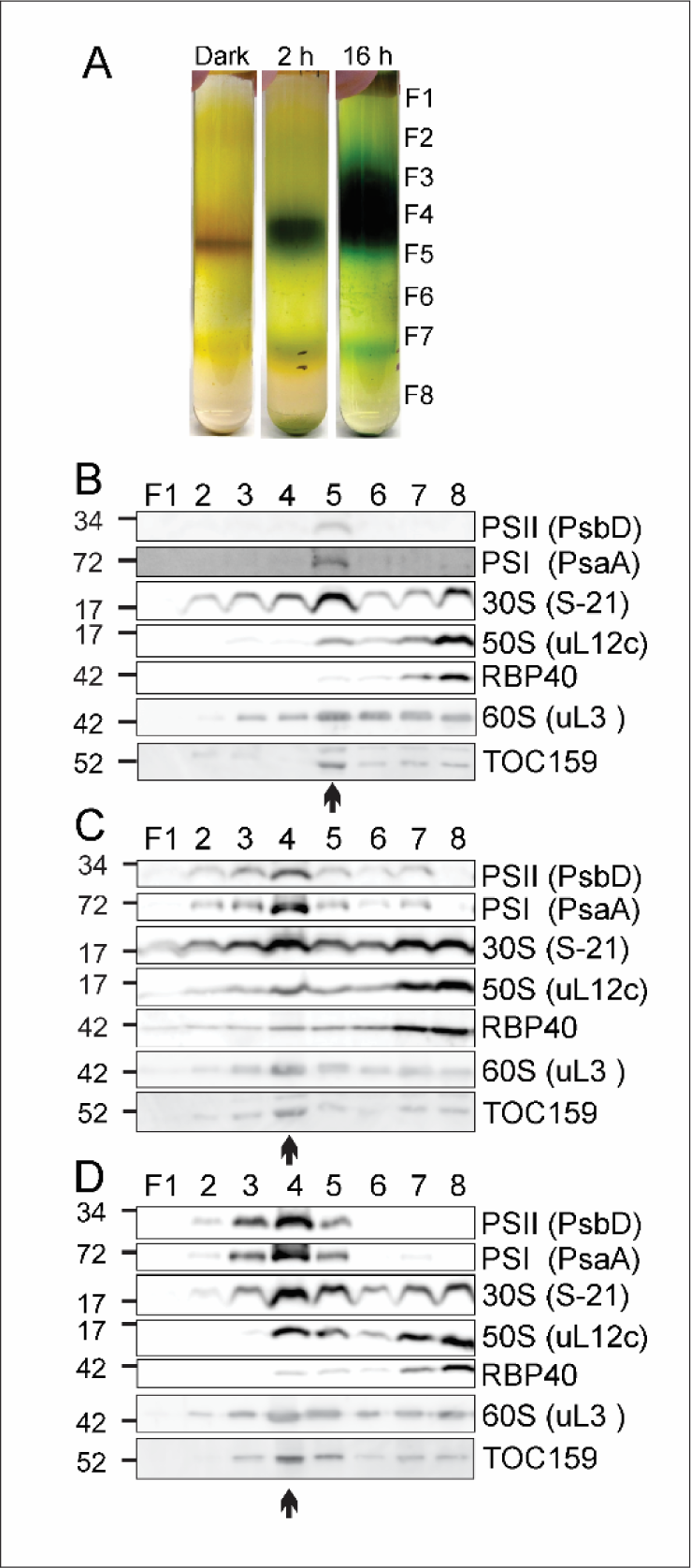
Density profiles of chloroplast membranes before, during and after TM biogenesis in *y1* greening. (A) Sucrose density gradient ultracentrifugation fractionated membranes from dark-*y1* cells, 2h-*y1* cells, and 16h-*y1* cells. During isopycnic ultracentrifugation, membranes floated from a dense sucrose cushion (F8) to equilibrate in the gradient by buoyant density and were then collected as fractions F1-F7. Immunoblot analyses of fractions from (B) dark-*y1* cells, (C) 2h-*y1* cells and (D) 16h-*y1* cells revealed levels of markers for the photosystems (PsbD and PsaA) and biogenesis: the chloroplast ribosome subunits of 30S (S-21) and 50S (uL12c), the translation activator RBP40, cytoplasmic ribosomes (uL3) and the TOC complex (TOC159). (B-D) arrows point to fractions with highest levels of cytoplasmic and chloroplast ribosome markers (see text). (B-D) Lanes were loaded with equal proportions of the total rather than by equal protein mass amounts. Molecular mass markers are in kDa.

### LDM has features of a platform for translation initiation

Our results support an early role of LDM in translation for photosystem biogenesis in the chloroplast. LDM was most prominent in dark-*y1* cells, correlating with their poise to activate translation during greening (Figure 1 B) (Sun et al., 2019). LDM preparations had a higher ratio of 30S to 50S chloroplast ribosomal subunits compared to TM or CTM (Figure 1, compare 30S and 50S markers in LDM fractions F2-F4 in B versus CTM fractions F5-F7 in B-D or the TM fraction F4 in C or D). This feature supports an early role because the initiation phase of translation involves the 30S subunit but not the 50S subunit (Zerges and Rochaix, 1998; Laursen et al., 2005). This LDM role in organizing translation initiation also is supported by the appearance of the 50S subunit marker protein to the low-density fractions during the first 2 hours of greening, e.g. for the assembly of the 70S ribosome (compare Figure 1 C, to B and D, respectively).

### Proteomic results support roles of LDM and CTM as platforms for chloroplast protein synthesis and assembly

We carried out proteomic analyses of LDM and CTM to obtain a comprehensive characterization of their functions. LDM from wild-type cells undergoing exponential growth and TM biogenesis was isolated using the published protocol, rather than from F1-F4 from dark-*y1* cells (Figure 1 B). This was to benefit from the purity achieved from the two sequential isopycnic sucrose gradient purification steps (Zerges and Rochaix, 1998). Immunoblot results confirmed that the LDM and TM preparations had the expected relative levels of markers for photosynthesis and biogenesis (Figure 2 A). TM had higher levels of PSI and PSII markers. In contrast, LDM had higher levels of markers for translation and chloroplast biogenesis: the chloroplast ribosome (S-21), a protein of the TOC chloroplast protein import translocon (Toc159), RBP40, and an enzyme in the chlorophyll branch of the tetrapyrrole biosynthetic pathway (PORA). We found that LDM preparations had higher levels than do TM of the AtpB subunit of the chloroplast ATP synthase, a complex believed to be only in TM (Hahn et al., 2018).

**Figure 2.**
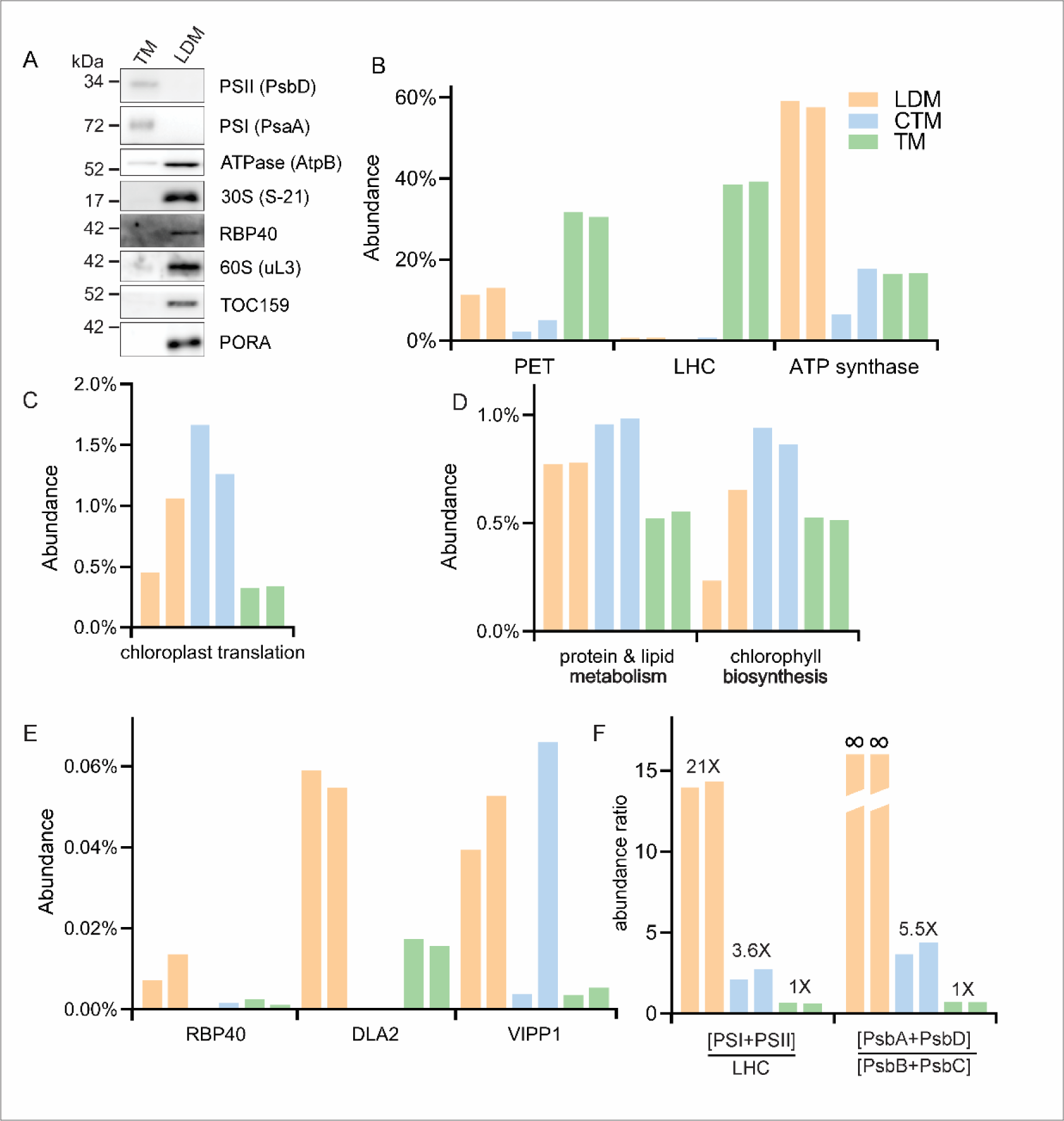
Proteomic analyses of LDM and CTM support their roles as translation platforms for TM biogenesis. (A) Immunoblot analyses revealed the relative levels in LDM and TM of marker proteins for PSII (PsbD), PSI (PsaA), the chloroplast ATP synthase (AtpB), the chloroplast translation (S-21 and RBP40), the cytoplasmic ribosome (uL3), chlorophyll biosynthesis (PORA), and chloroplast protein import (TOC159). Molecular mass markers are in kDa. (B-F) Bars are color-coded as follows: LDM, orange; CTM, blue; TM, green. Bar pairs are the biological replicate trials 1 and 2. (B-E) Bar heights indicate the representations by the functional categories, as the sum of the abundances of the proteins therein as the percent total abundances of all proteins in the sample. Categories were (B) photosynthetic complexes of PET, LHCs and ATP synthase, (C) chloroplast translation, (D) chloroplast protein and lipid metabolism, trafficking and assembly and chlorophyll biosynthesis (E) RBP40, RBP60/DLA2, VIPP1, and (F) abundance ratios of photosystem proteins to LHC proteins and PsbA and PsbD to PsbB and PsbC. Values above each bar pair represent fold difference relative to TM. The two left-most bars for LDM were assigned ∞ because these ratios cannot be calculated with PsbB and PsbC abundance values of zero in the denominator.

Our proteomic analyses of LDM included comparisons to TM with a quantitative approach based on their differential metabolic labelling with stable N isotopes (Gouw et al., 2010). To characterize CTM, we carried out label-free proteomic analysis of the sucrose density gradient fraction F5 of dark-*y1* cell membranes, which was validated by the results presented above and in Figure 1 B. Two independent biological replicates were carried out from separate cultures for each membrane type. In the LDM preparations, the numbers of annotated proteins identified were 192 and 260, respectively, whereas in the CTM preparations, they were 756 and 671 (table 1, table 2, Supplementary data file 1). Biogenic proteins were assorted into the following categories: 1) the routing and assembly of TM lipids and proteins, 2) the chlorophyll branch of tetrapyrrole biosynthesis, and 3) chloroplast translation, and 4) cytoplasmic translation. Photosynthetic proteins were categorized by their function in PET, LHCs, or ATP synthase. Proteins of the PET system were more numerous in TM (51) than in LDM (30) and CTM (27). The representations of the protein function categories in LDM, CTM and TM were compared by plotting the sum of protein abundance values in each as a percentage of the total protein abundance in the sample (Figure 2 B-E). Two biological replicate trials yielded similar results. For the few cases where the results of the trials differed, we tempered our interpretations of them accordingly (Discussion). PET and LHC proteins were most abundant in TM, their known location (Figure 2 B). These results provide evidence supporting these biogenic roles of LDM and CTM and confirmed the unexpectedly high abundance of ATP synthase in LDM.

**Table 1.**
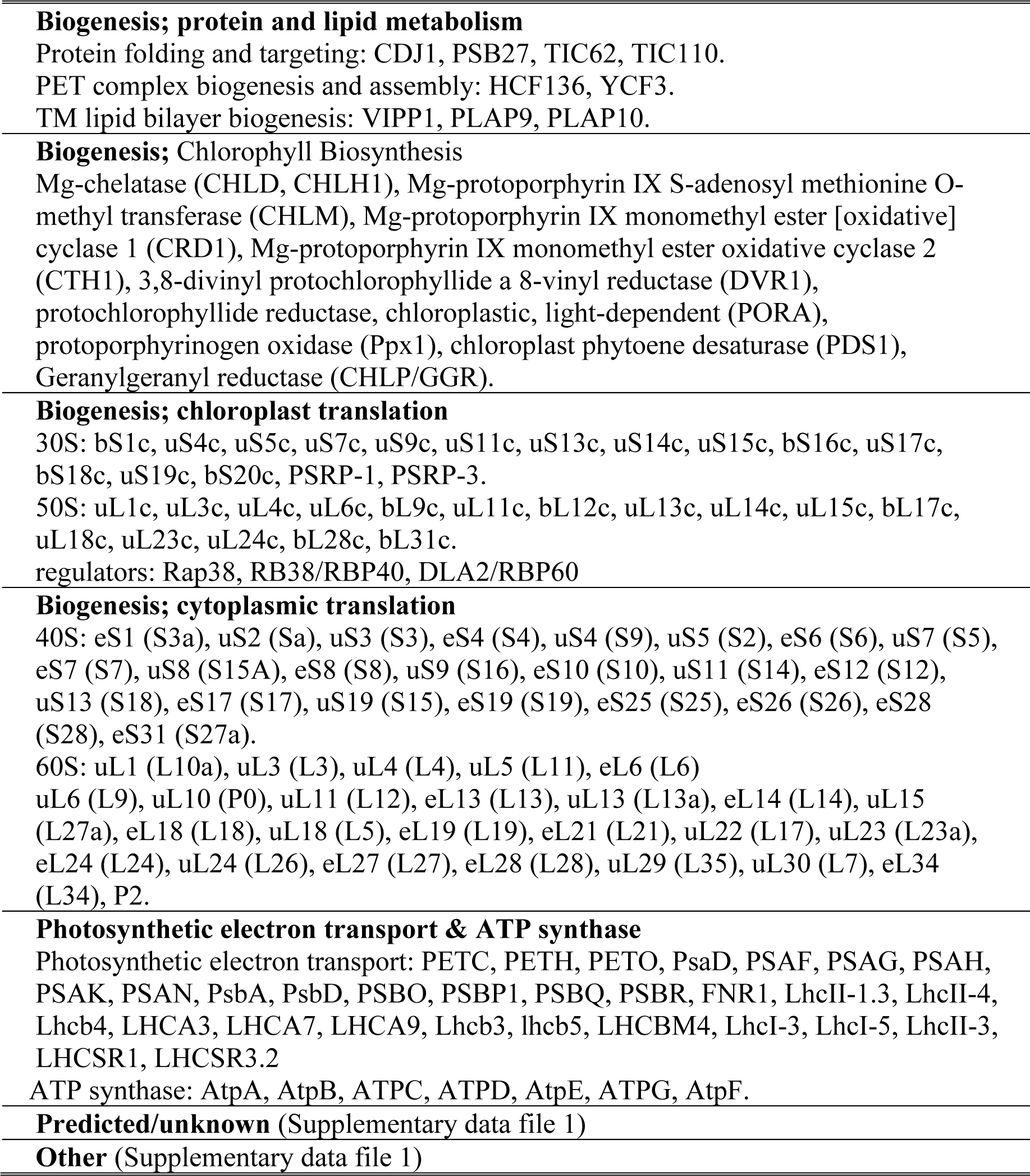
The LDM proteome. The proteins identified in LDM are listed according to their known functions in chloroplast biogenesis or photosynthesis. Ribosomal protein names are provided according to the revised nomenclature, with the previous names for cytoplasmic ribosomal proteins given in parentheses (Ban et al., 2014). The older names are not provided for chloroplast ribosomal proteins because the numbers therein did not change (Willmund et al., 2022). For additional information see Supplementary data file 1.

**Table 2.**
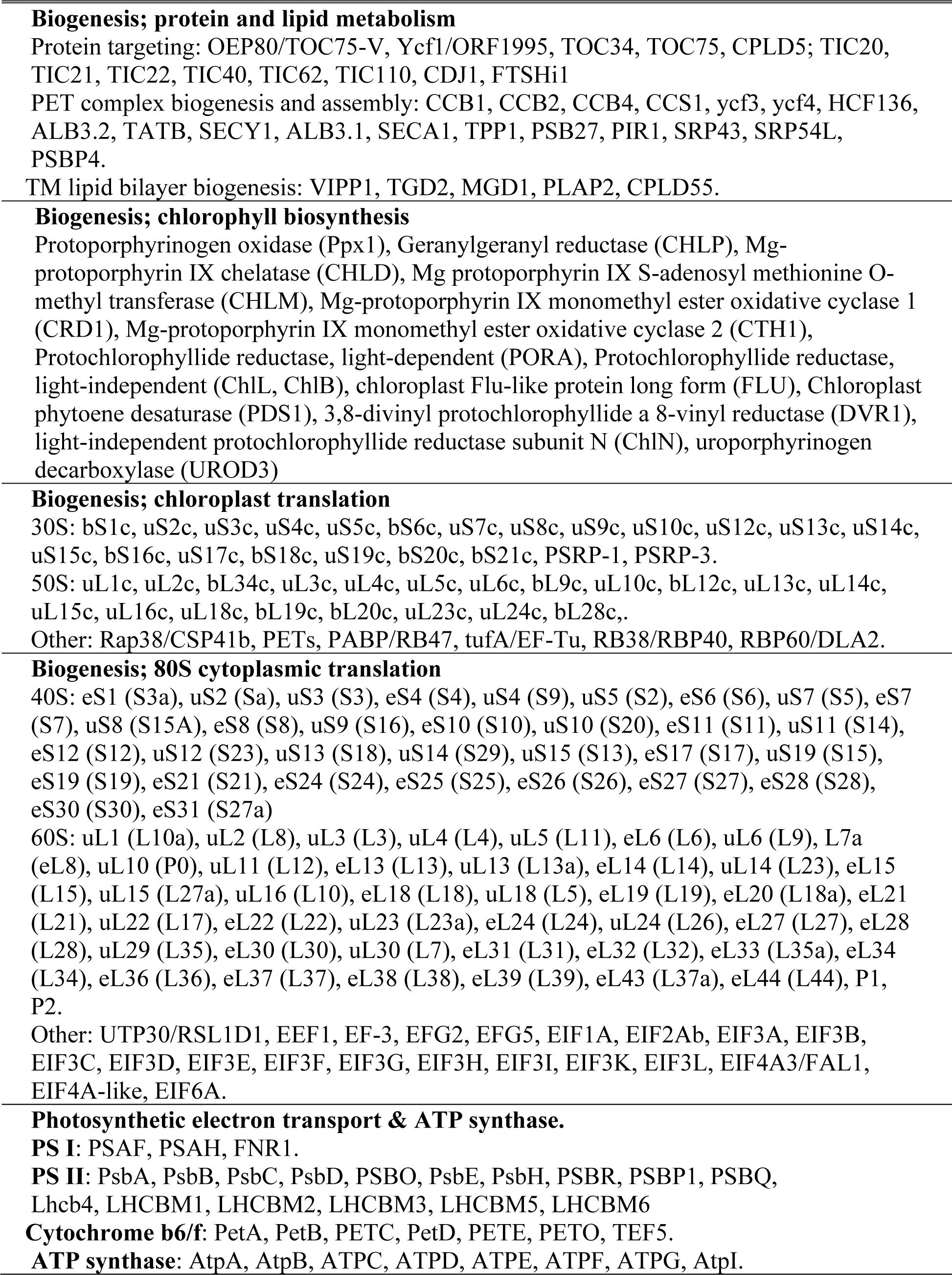
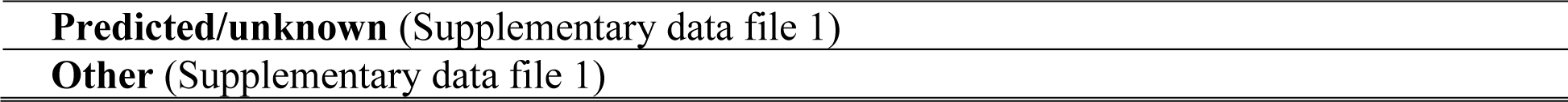
The CTM proteome. Ribosomal protein names are according to the revised nomenclature with the previous names for cytoplasmic ribosomal proteins given in parentheses (Ban et al., 2014). For additional information see Table 1 legend and Supplementary data file 1.

### Evidence of associations between membranes with cytoplasmic ribosomes and chloroplast ribosomes

Translation by chloroplast ribosomes in the T-zone is aligned with translation by cytoplasmic ribosomes on a domain of the chloroplast envelope (Sun et al., 2024). These chloroplast-bound cytoplasmic ribosomes synthesize proteins encoded by the nuclear genome for localized import into the chloroplast. This spatial alignment of the dual translation systems predicts connectivity between the membranous platforms in the respective compartments. Indeed, cytoplasmic ribosomal proteins were over 10-fold more abundant in CTM than in LDM or TM and this correlated with the highest levels of chloroplast ribosomal proteins (Figure 3 A). Similarly, envelope marker proteins were more abundant in CTM than in either LDM or TM (Figure 3 B). In our membrane fractionation experiments, we observed cofractionation of proteins of the cytoplasmic ribosome, the chloroplast ribosome, and the TOC translocon marker protein TOC159 (Figure 1 B-D, see arrows below each panel) (Schottkowski et al., 2012). Moreover, markers of translation in the chloroplast and cytoplasm together decreased in buoyant density during greening (Figure 1 B-D, F5 in dark-*y1* cells and F4 in 2h- and 16h-*y1* cells). Co-immunoprecipitation was observed between the cytoplasmic ribosomal marker (uL3) and the chloroplast ribosome (S-21) (Figure 3 C). Finally, in TEM images, we observed potential membranous connections between the translation systems; clusters of cytoplasmic ribosomes on the chloroplast envelope outside contact sites between the ends of thylakoid lamellae and the envelope in the T-zone (Figure 3 D). Similar contact sites, such as thylakoid convergence zones, have been proposed to be locations of TM biogenesis (Engel et al., 2015; Rast et al., 2019; Ostermeier et al., 2024).

**Figure 3.**
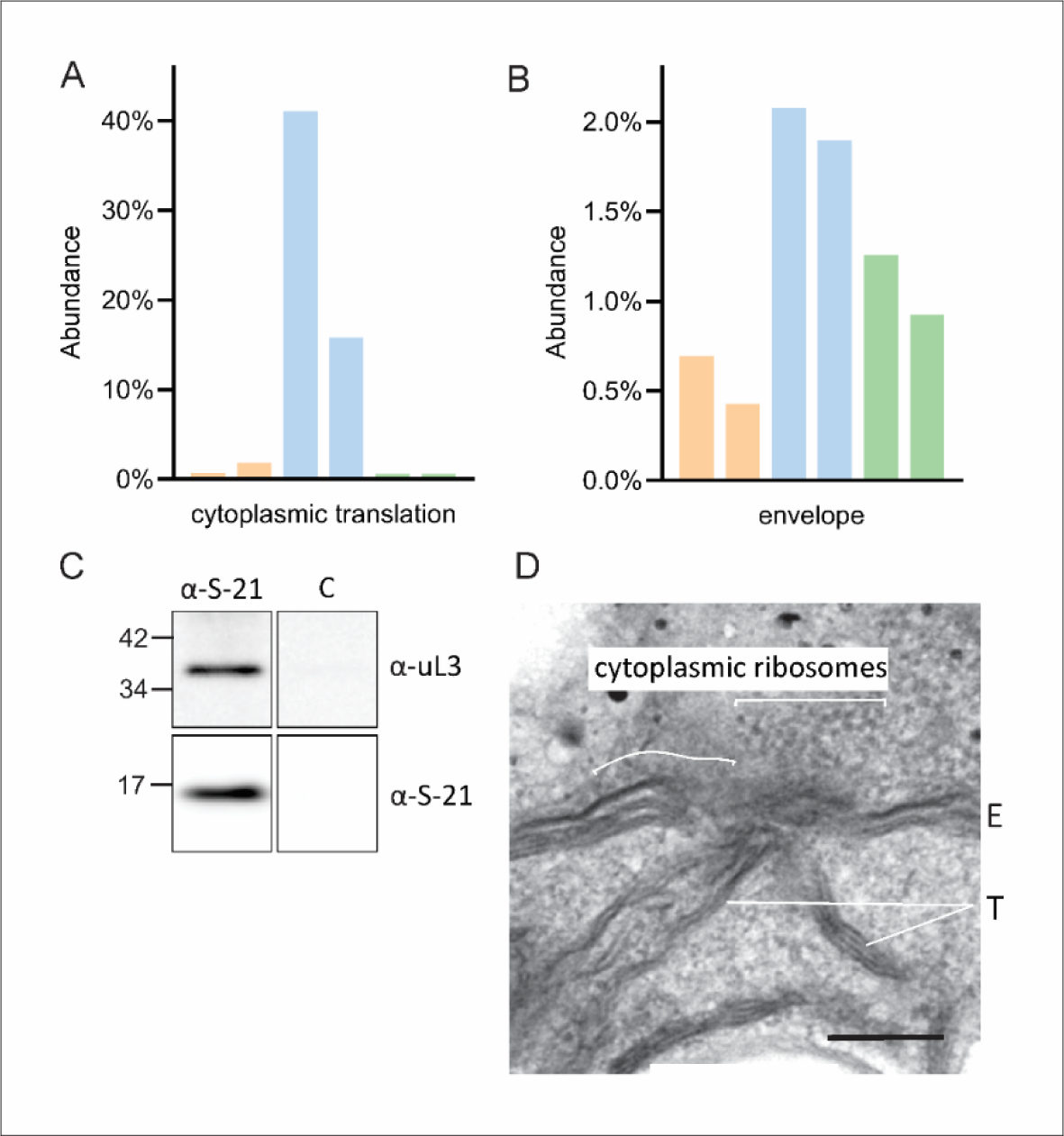
Evidence of physical connections between membranous platforms for cytoplasmic ribosomes and chloroplast ribosomes. (A and B) Abundance (y-axis) is the sum of protein abundances in the category as the precent of the total protein abundance in the sample. Each pair of bars represents results of the two independent biological replicate experiments. Trial 1 and trial 2 are the left and right bars, respectively. The protein function categories are A) cytoplasmic ribosomal proteins and (B) envelope proteins. (C) Membranes with the cytoplasmic ribosome marker protein uL3 co-immunoprecipitated with membranes with the chloroplast ribosome marker S-21. In the negative control (“C”), antibody against S-21 was excluded. Molecular mass markers are in kDa. (D) A TEM image shows apparent contacts between the chloroplast envelope and the termini of thylakoid lamellae (T) in the T-zone. Cytoplasmic ribosomes are seen in clusters near and on the outer membrane of the chloroplast envelope (E). bar = 200 nm.

## DISCUSSION

Membrane domains compartmentalize biogenic processes in organelles and bacteria (Hoffman et al., 2019; Rast et al., 2019; Zorkau et al., 2021; Mahbub and Mullineaux, 2023). LDM and CTM roles as membranous platforms for translation and photosystem assembly are supported by their enrichment for proteins in these processes and their distinction from TM. For example, LDM and CTM were enriched in proteins of the chloroplast ribosome, protein and lipid metabolism and trafficking, and chlorophyll biosynthesis (Figure 2 C and D). The identification of chlorophyll biosynthetic enzymes supports these roles because this photopigment binds to apo-proteins during or immediately following their synthesis for apoprotein stabilization, functionality and to prevent phototoxicity by free chlorophyll. The minor amounts of photosystem subunits identified are consistent with their synthesis and assembly at LDM and CTM. Such evidence in cyanobacteria supports the plasma membrane as the location of photosystem assembly (Figure 2 B) (Smith and Howe, 1993; Zak et al., 2001; Stengel et al., 2012). The detection of chloroplast ribosomes and assembly factors in TM could reflect their roles in photosystem repair (Chotewutmontri et al., 2020), contamination of our TM preparations by LDM or CTM, or the occurrence of some de novo photosynthesis complex biogenesis in TM.

A spatial-temporal pathway or “assembly line” of TM biogenesis is supported by our results (Figure 4) (Schottkowski et al., 2012). An early role of LDM is supported by its higher abundance in *y1*-dark cells and their poise to activate high levels of translation by chloroplast ribosomes than in 2h-*y1* or 16h-*y1* cells (Figure 1 B, C, and D) (Malnoe et al., 1988; Sun et al., 2019). LDM preparations had chloroplast ribosomal proteins and were enriched in the translational activators RBP40 and RBP60/DLA2, relative to CTM and TM (Figure 2 C and E). LDM from dark-*y1* cells had an elevated ratio of 30S to 50S chloroplast ribosomal subunits relative to CTM and TM, consistent with a role in the initiation phase of translation (Figure 1 B, Results). A protein that functions very early in the biogenesis of the TM lipid bilayer, VIPP1, was more abundant in LDM than in the other membrane types (Figure 2 E) (Gupta et al., 2021). The early role of LDM in photosystem biogenesis is further supported by the relative levels of different photosynthesis complexes and specific subunits. LDM had approximately a 20-fold higher abundance ratio of photosystem proteins to LHC proteins compared to TM (Figure 2 F). This result aligns with the assembly of photosystems in LDM before the LHCs join them later in CTM (McCormac and Greenberg, 1992; Dreyfuss and Thornber, 1994; Minagawa and Takahashi, 2004; Mehra et al., 2024). The detection of PSII core subunits PsbA and PsbD but not PsbB or PsbC in LDM supports this early role in PSII core assembly, because the former are synthesized and assembled before the latter (Choquet and Wollman, 2023) (Figure 2 F, Supplementary Data file 1). Finally, the high abundance of chloroplast ATP synthase proteins in LDM could reflect its biogenesis in LDM, which is known to occur before photosystem biogenesis (Figure 2 A and B) (Zones et al., 2015; Sun et al., 2019). Alternatively, ATP synthase in LDM could have a moonlighting function in TM biogenesis, analogous to the role of mitochondrial ATP synthase in the curvature of the inner membrane to form cristae (Blum et al., 2019).

**Figure 4.**
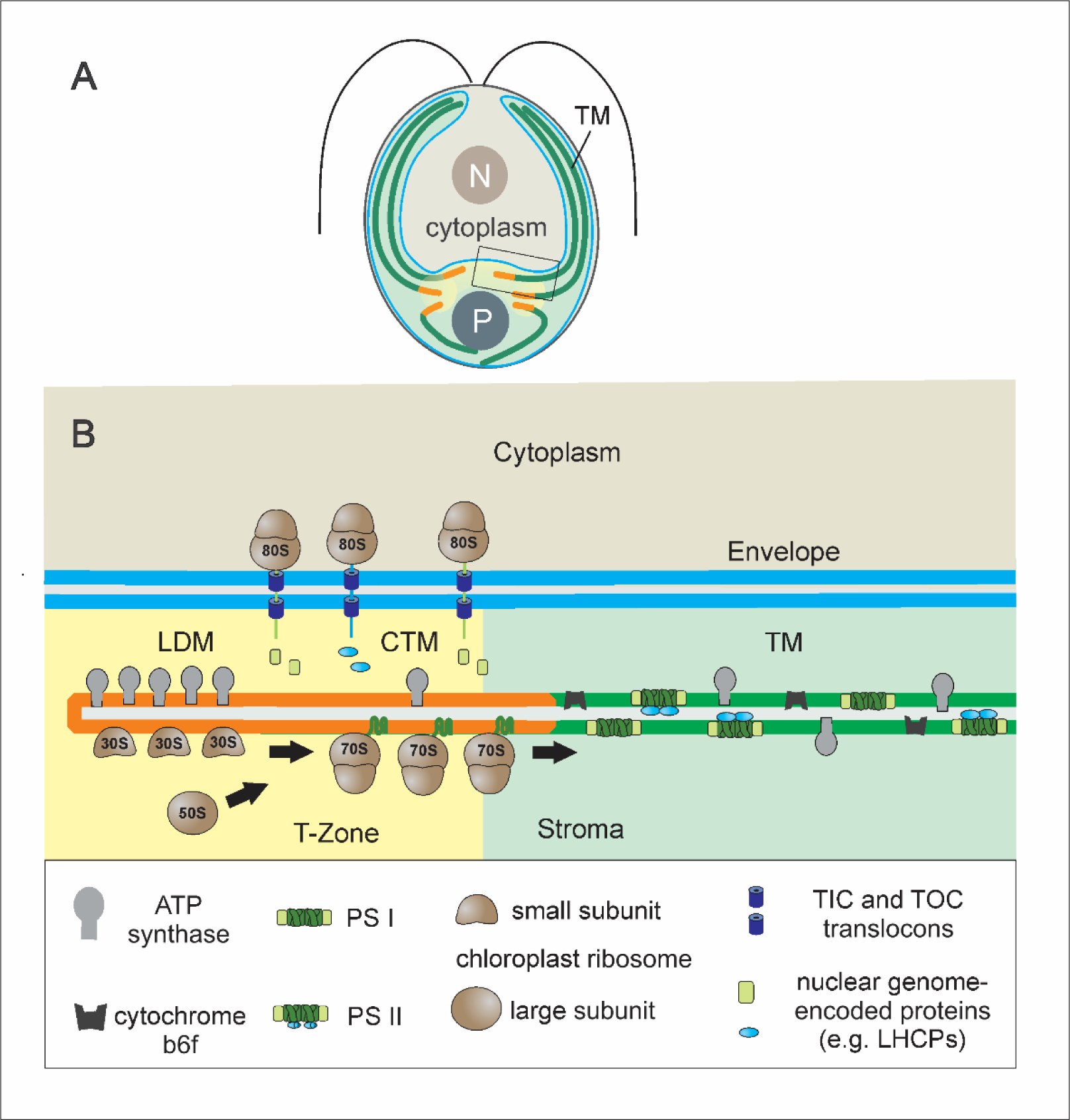
The model for the organization of LDM and CTM in the chloroplast. (A) An illustration of a *Chlamydomonas* cell shows the nucleus (N), cytoplasm, and chloroplast (light green) with its T-zone (yellow), pyrenoid (P), thylakoid vesicles (dark green, TM), envelope (blue), and LDM and CTM (orange). The rectangular box shows the region illustrated in B. (B) LDM is a platform for preliminary stages of translation and photosystem biogenesis. On CTM, 70S ribosomes synthesize chloroplastic genome-encoded subunits of PSI and PSII for assembly. The resulting PSI and PSII reaction centers are bound by subunits expressed from the nuclear genome and imported by the TOC and TIC translocons (purple cylinders) (Schottkowski et al., 2012). Newly assembled photosystems diffuse from CTM to TM throughout the chloroplast.

The role of CTM as a platform for ongoing translation and photosystem assembly is supported by its prominence in 2h-*y1* cells, thus correlating with the highest rates of translation for photosystem biogenesis of the three stages analyzed (Figure 1 B-D) (Sun et al., 2019). CTM also had the highest abundances of proteins of the chloroplast ribosome, the metabolism and trafficking of chloroplast proteins and lipids, and chlorophyll biosynthesis and higher abundance ratios of photosystem proteins to LHC proteins and of PsbA and PsbD to PsbB and PsbC, compared to TM (Figure 2 C-E) (see previous paragraph). These ratios are intermediate between those of LDM and TM, and thus support the proposed position of CTM between LDM and TM (Figure 4 B).

We posit that LDM gives rise to CTM as it increases in buoyant density due to the assembly of 70S chloroplast ribosomes and newly synthesized TM proteins. Accordingly, the 50S subunit and RBP40 appeared in the LDM fractions during the initial 2 h of *y1* greening (Figure 1 B and C). Assembling photosystems diffuse from LDM to CTM in a contiguous membrane where they assemble with LHCs, whereupon the newly assembled photosystem-LHC supercomplexes undergo further outward diffusion to populate the membranes of thylakoid lamellae throughout the chloroplast (Figure 4). Directionality is imposed by localized assembly in LDM and CTM. The rate of lateral diffusion of complexes in TM (∼0.5 µm/min, Kirchhoff et al., 2008) is sufficient for photosystems generated in the T-zone to populate the entire thylakoid network by diffusion over several microns in the ∼5-h window of photosystem biogenesis in the cell cycle (Zones et al., 2015; Sun et al., 2019).

Does our model account for the published evidence supporting the long-standing belief that PET complex biogenesis occurs at stroma-exposed TM? The electron microscopy evidence of ribosome-bound “rough” thylakoids does not contradict our model because chloroplast ribosomes were seen primarily near the termini of thylakoid lamella in cell lysates (Figure 4) (Chua et al., 1976; Yamamoto et al., 1981). The biochemical analyses of ribosome-bound TM used crude pellet fractions from differential centrifugation, which could have contained LDM and CTM as well as free ribosomes and polysomes (Falk, 1969; Chua et al., 1976; Yamamoto et al., 1981; Jagendorf and Michaels, 1990; Zoschke and Barkan, 2015). Our model does not challenge the occurrence of translation on stroma-exposed TM throughout the chloroplast, the likely location of *psbA* translation for the PSII damage-repair cycle (Zoschke and Bock, 2018). The evidence that the chloroplast envelope is the location of biosynthesis of the galactolipids of the TM bilayer and the photopigments (chlorophyll and carotenoids) of the PET complexes and LHCs is consistent with the localization of these processes at LDM and CTM. These biochemical studies analyzed purified envelope which could have contained LDM, due to their overlapping buoyant densities (Zerges and Rochaix, 1998). There is little microscopy evidence of envelope localization of chlorophyll and galactolipid biosynthesis in plant chloroplasts, whereas fluorescence microscopy revealed chlorophyll biosynthesis localized to the T-zone in Chlamydomonas (Sun et al., 2019). Our results and model offer an intriguing answer to the long-standing question regarding how newly synthesized chlorophyll and galactolipids undergo bulk transport from the envelope to thylakoids: they don’t. Rather, they are biosynthesized and assembled in LDM and CTM to form PET complexes and the membrane bilayer and then routed to TM as described above (Figure 4 B).

Spatial coordination of the cytoplasmic and chloroplast translation systems was demonstrated recently and proposed to localize newly synthesized proteins encoded by the nuclear and plastid genomes to the T-zone for photosystem assembly (Sun et al., 2024). Our results suggest that this coordination involves a physical association of membranous translation platforms for chloroplast ribosomes and cytoplasmic ribosomes (Figure 3).

Localized translation on specialized membranes has also been reported in the other semiautonomous organelle, mitochondria, and the cyanobacterial evolutionary relatives of the plastids (Zak et al., 2001; Stengel et al., 2012; Rast et al., 2019; Mahbub and Mullineaux, 2023). For example, in yeast mitochondria, domains of the inner membrane organize translation by mitochondrial ribosomes for the biogenesis of the respiratory electron transport system and ATP synthase in a spatiotemporal biogenic pathway (Vogel et al., 2006; Garcia et al., 2007; Stoldt et al., 2018). Hence, assembly line models are emerging in the biogenesis of bioenergetic membranes in semiautonomous organelles (Formosa and Ryan, 2018).

## EXPERIMENTAL PROCEDURES

### Strains and culture conditions

All cultures were grown at 24 °C under a light intensity of 150 µEꞏm^-2^ꞏs^-1^ and with orbital shaking unless stated otherwise. Cells were collected by centrifugation (3,000 g, 5 min at RT). LDM was isolated from CC-400 (CW15, *MT*+) cultured in HSM with 1.0% (w/v) sorbitol. TEM was performed on the wild-type strain CC-125 cultured to 1×10^6^ cellsꞏml^-1^ in high salt minimal (HSM) medium (Harris, 1989) in a 12h light:12 h dark regime as described previously (Sun et al., 2019). Cells were harvested at the fourth hour of the light cycle. For differential quantitative MS using metabolic labeling in the analysis of LDM vs TM, cultures were grown on TAP medium containing either [^14^N]NH_4_Cl (LDM) or [^15^N]NH_4_Cl (TM). *y1* (CC-1168) was cultured on TAP medium in the dark and greening was induced by exposure to a white light at 150 µEꞏm^-2^ꞏs^-1^. Strains were obtained from the *Chlamydomonas* Resource Center (https://www.chlamycollection.org).

### Membrane fractionation

*y1* membranes for the density profiles in Figure 1 and the proteomic analysis of CTM and TM were prepared as described previously (Schottkowski et al., 2012). The MS analyses were carried out on LDM purified as reported previously (Zerges and Rochaix, 1998).

### Immunoblot Analysis

For the immunoblots in Fig 1 B-D, equal proportions of the fractions from the same number of cells in each gradient were resuspended in SDS-PAGE loading buffer and resolved by SDS-PAGE (12% acrylamide) (Sambrook and Russell, 2001). Proteins were transferred to PVDF (BIO-RAD) and reacted with primary and secondary antibodies as described previously (Sambrook and Russell, 2001). The primary antibodies and the dilutions were: α-TOC159 (1:8,000) (Agrisera, AS07-239), α-AtpB (1:6,000), α-S-21 (1:3,000) (Randolph-Anderson et al., 1989), α-uL12c (1:10,000), α-uL3 (1:6,000), α-PsbD (1:4,000) (Agrisera, AS06-146), α-PsaA (1:60,000) (Prof. Kevin Redding, Arizona State University), α-RBP40 (1:1,000) (Prof. Jorg Nickelsen, Ludwig Maximilian University) and α-PORA (1:20,000) (Prof. Jürgen Soll, Ludwig Maximilian University) (Supplemental Materials). The secondary antibody was horseradish peroxidase-conjugated goat anti-rabbit IgG antibody (KPL). Signals were detected using an ECL substrate (Thermo-Fisher) with an Imager 600 (Amersham/GE) according to the manufacturer’s protocols.

### Mass spectrometry (see Supplementary Materials)

#### Immunoprecipitation

Chloroplasts were isolated from CC-400 (CW15, MT+) cultured in HSM with 1.0 % (w/v) sorbitol, as described (Sun et al., 2024). Chloroplasts were collected from the 45-65% interface, diluted with 4 vol of ice cold IB buffer [300 mM Sorbitol, 50 mM HEPES-KOH pH 7.5, 25 mM MgCl_2_, 0.1% (w/v) BSA], pelleted by centrifugation (670 x g, 1 min, 4°C). The chloroplast pellet was resuspended in ice cold TKM buffer [25 mM MgCl_2_, 20 mM KCl, 10 mM Tricine-HCl pH 7.5, protease inhibitor cocktail for plants (Sigma-Aldrich)]. Chloroplasts were broken by four passes through an ice-chilled French Pressure Cell at 1,000 psi. The lysate was ultracentrifuged at 100,000 x g for 1 h at 4°C. The pellet was resuspended in ice cold TKM buffer as input. α-S-21 (1:1,000) was incubated with the input overnight at 4 °C with slow rotation. In the control, pre-immune serum (1:1,000) was incubated with the input overnight at 4 °C with slow rotation. DynabeadsTM Protein A (ThermoFisher) was prepared according to the manufacturer’s protocol. The prepared beads were then incubated with the sample or the control mixture at 4 °C for 2 hours with slow rotation. The beads were washed according to the manufacturer’s protocol. The beads were then resuspended in SDS-PAGE loading buffer and incubated at 70 °C for 5 min.

### Accession Numbers

Sequence data from this article can be found in the GenBank/EMBL data libraries under accession numbers provided in Supplementary data file 1.

## SUPPLEMENTARY DATA

Date file S1. Proteomic results

Supplementary information and methods

## ACKNOWLEDGMENTS

For infrastructure and technical support, we thank the Centre for Structural & Functional Genomics (Concordia University), the Centre for Biological Applications of Mass Spectrometry (Concordia University) and Jeannie Mui and the Facility for Electron Microscopy Research (McGill University). For generous gifts of antibodies, we thank Dr. Elizabeth Harris (αAtpB and α-uL3, Duke University), Prof. Jorg Nickelsen (αRBP40), Dr. Kevin Redding (αPsaA) and Dr. Katrin Philippar and Prof. Jurgen Soll (α-PORA) (Ludwig Maximilian University).

## AUTHOR CONTRIBUTIONS

YS, SB, and WZ designed the research; YS, SB, MV-P performed the research; YS, SB, HJ and WZ analyzed data; YS and WZ wrote the paper.

## SHORT LEGENDS FOR SUPPORTING INFORMATION

Supplementary material: Background information regarding antibodies against the ribosomal proteins. Detailed methodological information for the sample preparation, analyses and processing are provided for the mass spectrometry and proteomics.

Supplemental Data File 1. The proteomic results are presented.

## DATA AVAILABILITY

Data are available upon reasonable request. The raw MS data will be available on PRIDE.

## FUNDING AND ADDITIONAL INFORMATION

This work was supported by NSERC Discovery Grant 217566 (WZ).

## REFERENCES

Ban N, Beckmann R, Cate JH, Dinman JD, Dragon F, Ellis SR, Lafontaine DL, Lindahl L, Liljas A, Lipton JM, et al (2014) A new system for naming ribosomal proteins. Curr Opin Struct Biol 24: 165–9

Billakurthi K, Loudya N (2024) Co-translational import of nuclear-encoded proteins into the chloroplast in Chlamydomonas reinhardtii. Plant Physiology kiae310

Blum TB, Hahn A, Meier T, Davies KM, Kühlbrandt W (2019) Dimers of mitochondrial ATP synthase induce membrane curvature and self-assemble into rows. Proceedings of the National Academy of Sciences 116: 4250–4255

Cahoon AB, Timko MP (2000) yellow-in-the-dark mutants of Chlamydomonas lack the CHLL subunit of light-independent protochlorophyllide reductase. Plant Cell 12: 559–68.

Choquet Y, Wollman F-A (2023) Chapter 19 - The assembly of photosynthetic proteins. In AR Grossman, F-A Wollman, eds, The Chlamydomonas Sourcebook (Third Edition). Academic Press, London, pp 615–646

Chotewutmontri P, Barkan A (2016) Dynamics of Chloroplast Translation during Chloroplast Differentiation in Maize. PLoS Genet 12: e1006106

Chotewutmontri P, Williams-Carrier R, Barkan A (2020) Exploring the Link between Photosystem II Assembly and Translation of the Chloroplast psbA mRNA. Plants 9: 152

Chua NH, Blobel G, Siekevitz P, Palade GE (1976) Periodic variations in the ratio of free to thylakoid-bound chloroplast ribosomes during the cell cycle of Chlamydomonas reinhardtii. J Cell Biol 71: 497–514

Dreyfuss BW, Thornber JP (1994) Organization of the Light-Harvesting Complex of Photosystem I and Its Assembly during Plastid Development. Plant Physiology 106: 841– 848

Engel BD, Schaffer M, Kuhn Cuellar L, Villa E, Plitzko JM, Baumeister W (2015) Native architecture of the Chlamydomonas chloroplast revealed by in situ cryo-electron tomography. Elife. doi: 10.7554/eLife.04889

Falk H (1969) ROUGH THYLAKOIDS: POLYSOMES ATTACHED TO CHLOROPLAST MEMBRANES 10.1083/jcb.42.2.582. J Cell Biol 42: 582–587

Formosa LE, Ryan MT (2018) Mitochondrial OXPHOS complex assembly lines. Nat Cell Biol 20: 511–513

Garcia M, Darzacq X, Delaveau T, Jourdren L, Singer RH, Jacq C (2007) Mitochondria-associated Yeast mRNAs and the Biogenesis of Molecular Complexes. MBoC 18: 362– 368

Gouw JW, Krijgsveld J, Heck AJ (2010) Quantitative proteomics by metabolic labeling of model organisms. Mol Cell Proteomics 9: 11–24

Gupta TK, Klumpe S, Gries K, Heinz S, Wietrzynski W, Ohnishi N, Niemeyer J, Spaniol B, Schaffer M, Rast A, et al (2021) Structural basis for VIPP1 oligomerization and maintenance of thylakoid membrane integrity. Cell 184: 3643–3659.e23

Hahn A, Vonck J, Mills DJ, Meier T, Kühlbrandt W (2018) Structure, mechanism, and regulation of the chloroplast ATP synthase. Science 360: eaat4318

Harris EH (1989) The Chlamydomonas sourcebook: a comprehensive guide to biology and laboratory use. Academic Press, San Diego

Hoffman AM, Chen Q, Zheng T, Nicchitta CV (2019) Heterogeneous translational landscape of the endoplasmic reticulum revealed by ribosome proximity labeling and transcriptome analysis. J Biol Chem 294: 8942–8958

Jagendorf AT, Michaels A (1990) Rough thylakoids: translation on photosynthetic membranes. Plant Science 71: 137–145

Järvi S, Suorsa M, Aro E-M (2015) Photosystem II repair in plant chloroplasts — Regulation, assisting proteins and shared components with photosystem II biogenesis. Biochimica et Biophysica Acta (BBA) - Bioenergetics 1847: 900–909

Kettunen R, Pursiheimo S, Rintamaki E, Van Wijk KJ, Aro EM (1997) Transcriptional and translational adjustments of psbA gene expression in mature chloroplasts during photoinhibition and subsequent repair of photosystem II. Eur J Biochem 247: 441–8.

Kirchhoff H, Haferkamp S, Allen JF, Epstein DBA, Mullineaux CW (2008) Protein Diffusion and Macromolecular Crowding in Thylakoid Membranes. Plant Physiology 146: 1571–1578

Klein RR, Mason HS, Mullet JE (1988) Light-regulated translation of chloroplast proteins. I. Transcripts of psaA-psaB, psbA, and rbcL are associated with polysomes in dark-grown and illuminated barley seedlings. J Cell Biol 106: 289–301.

Laursen BS, Sorensen HP, Mortensen KK, Sperling-Petersen HU (2005) Initiation of protein synthesis in bacteria. Microbiol Mol Biol Rev 69: 101–23

Mahbub M, Mullineaux CW (2023) Locations of membrane protein production in a cyanobacterium. Journal of Bacteriology 205: e00209–23

Malnoe P, Mayfield SP, Rochaix JD (1988) Comparative analysis of the biogenesis of photosystem II in the wild-type and Y-1 mutant of Chlamydomonas reinhardtii. J Cell Biol 106: 609–16.

McCormac DJ, Greenberg BM (1992) Differential Synthesis of Photosystem Cores and Light-Harvesting Antenna during Proplastid to Chloroplast Development in Spirodela oligorrhiza 1. Plant Physiology 98: 1011–1019

Mehra HS, Wang X, Russell BP, Kulkarni N, Ferrari N, Larson B, Vinyard DJ (2024) Assembly and Repair of Photosystem II in Chlamydomonas reinhardtii. Plants 13: 811

Minagawa J, Takahashi Y (2004) Structure, function and assembly of Photosystem II and its light-harvesting proteins. Photosynth Res 82: 241–63

Nilsson R, Brunner J, Hoffman NE, van Wijk KJ (1999) Interactions of ribosome nascent chain complexes of the chloroplast-encoded D1 thylakoid membrane protein with cpSRP54. Embo J 18: 733–42

Ohad I, Siekevitz P, Palade GE (1967) Biogenesis of chloroplast membranes. II. Plastid differentiation during greening of a dark-grown algal mutant (Chlamydomonas reinhardi). J Cell Biol 35: 553–84.

Ostermeier M, Garibay-Hernández A, Holzer VJC, Schroda M, Nickelsen J (2024) Structure, biogenesis, and evolution of thylakoid membranes. The Plant Cell koae102

Randolph-Anderson BL, Gillham NW, Boynton JE (1989) Electrophoretic and immunological comparisons of chloroplast and prokaryotic ribosomal proteins reveal that certain families of large subunit proteins are evolutionarily conserved. J Mol Evol 29: 68–88

Rast A, Schaffer M, Albert S, Wan W, Pfeffer S, Beck F, Plitzko JM, Nickelsen J, Engel BD (2019) Biogenic regions of cyanobacterial thylakoids form contact sites with the plasma membrane. Nature Plants 5: 436–446

Rochaix J-D (2023) Historical Overview. The Chlamydomonas Sourcebook (Third Edition), 3rd ed. Academic Press, pp 1–22

Sambrook J, Russell DW (2001) Molecular cloning: a laboratory manual, 3rd ed. Cold Spring Harbor Laboratory Press, Cold Spring Harbor, N.Y.

Schottkowski M, Peters M, Zhan Y, Rifai O, Zhang Y, Zerges W (2012) Biogenic membranes of the chloroplast in Chlamydomonas reinhardtii. Proc Natl Acad Sci U S A 109: 19286–91

Schwarz C, Elles I, Kortmann J, Piotrowski M, Nickelsen J (2007) Synthesis of the D2 protein of photosystem II in Chlamydomonas is controlled by a high molecular mass complex containing the RNA stabilization factor Nac2 and the translational activator RBP40. Plant Cell 19: 3627–39

Smith D, Howe CJ (1993) The distribution of Photosystem I and Photosystem II polypeptides between the cytoplasmic and thylakoid membranes of cyanobacteria. FEMS Microbiology Letters 110: 341–347

Stengel A, Gugel IL, Hilger D, Rengstl B, Jung H, Nickelsen J (2012) Initial Steps of Photosystem II de Novo Assembly and Preloading with Manganese Take Place in Biogenesis Centers in Synechocystis. Plant Cell. doi: 10.1105/tpc.111.093914

Stoldt S, Wenzel D, Kehrein K, Riedel D, Ott M, Jakobs S (2018) Spatial orchestration of mitochondrial translation and OXPHOS complex assembly. Nat Cell Biol 20: 528–534

Sun Y, Bakhtiari S, Valente-Paterno M, Wu Y, Nishimura Y, Shen W, Law C, Dhaliwal J, Dai D, Bui KH, et al (2024) Chloroplast biogenesis involves spatial coordination of nuclear and organellar gene expression in Chlamydomonas. Plant Physiology kiae256

Sun Y, Valente-Paterno MI, Bakhtiari S, Law C, Zhan Y, Zerges W (2019) Photosystem Biogenesis Is Localized to the Translation Zone in the Chloroplast of Chlamydomonas. Plant Cell. doi: 10.1105/tpc.19.00263

Sun Y, Zerges W (2015) Translational regulation in chloroplasts for development and homeostasis. Biochim Biophys Acta 1847: 809–20

Uniacke J, Zerges W (2007) Photosystem II assembly and repair are differentially localized in Chlamydomonas. Plant Cell 19: 3640–54

Vogel F, Bornhovd C, Neupert W, Reichert AS (2006) Dynamic subcompartmentalization of the mitochondrial inner membrane. J Cell Biol 175: 237–47

Walter B, Hristou A, Nowaczyk MM, Schünemann D (2015) In vitro reconstitution of co-translational D1 insertion reveals a role of the cpSec–Alb3 translocase and Vipp1 in photosystem II biogenesis. Biochemical Journal 468: 315–324

Willmund F, Hauser C, Zerges W (2022) Translation and Protein Synthesis in the chloroplast. The Chlamydomonas Sourcebook 2:

Yamamoto T, Burke J, Autz G, Jagendorf AT (1981) Bound Ribosomes of Pea Chloroplast Thylakoid Membranes: Location and Release in Vitro by High Salt, Puromycin, and RNase. Plant Physiol 67: 940–949

Zak E, Norling B, Maitra R, Huang F, Andersson B, Pakrasi HB (2001) The initial steps of biogenesis of cyanobacterial photosystems occur in plasma membranes. Proc Natl Acad Sci U S A 98: 13443–8.

Zerges W, Rochaix J-D (1998) Low Density Membranes Are Associated with RNA-binding Proteins and Thylakoids in the Chloroplast of Chlamydomonas reinhardtii. J Cell Biol 140: 101–110

Zones JM, Blaby IK, Merchant SS, Umen JG (2015) High-Resolution Profiling of a Synchronized Diurnal Transcriptome from Chlamydomonas reinhardtii Reveals Continuous Cell and Metabolic Differentiation. Plant Cell 27: 2743–69

Zorkau M, Albus CA, Berlinguer-Palmini R, Chrzanowska-Lightowlers ZMA, Lightowlers RN (2021) High-resolution imaging reveals compartmentalization of mitochondrial protein synthesis in cultured human cells. PNAS. doi: 10.1073/pnas.2008778118

Zoschke R, Barkan A (2015) Genome-wide analysis of thylakoid-bound ribosomes in maize reveals principles of cotranslational targeting to the thylakoid membrane. Proc Natl Acad Sci U S A 112: E1678–87

Zoschke R, Bock R (2018) Chloroplast Translation: Structural and Functional Organization, Operational Control, and Regulation. Plant Cell 30: 745–770

## SUPPORTING INFORMATION REFERENCES

1 Schmidt, R. J., Myers, A. M., Gillham, N. W., and Boynton, J. E. (1984) Immunological similarities between specific chloroplast ribosomal proteins from Chlamydomonas reinhardtii and ribosomal proteins from Escherichia coli. Mol Biol Evol. 1, 317–34

2 Randolph-Anderson, B. L., Gillham, N. W., and Boynton, J. E. (1989) Electrophoretic and immunological comparisons of chloroplast and prokaryotic ribosomal proteins reveal that certain families of large subunit proteins are evolutionarily conserved. J Mol Evol. 29, 68–88

3 Banerjee, A. K., M, S., M, N., and Murty, U. S. (2010) Classification and clustering analysis of pyruvate dehydrogenase enzyme based on their physicochemical properties. Bioinformation. 4, 456–62

