## Supplemental Information for "Membranous translation platforms in the chloroplast of *Chlamydomonas reinhardtii*"

**Additional information regarding the antibodies.**

The antibodies against *Chlamydomonas* ribosomal proteins were raised against those proteins from highly purified ribosomal subunits in the laboratory of Drs. Gilham and Boynton and generously provided by Dr. Elisabeth Harris (Duke University). α-uL12c was raised against "L-30", the name in the nomenclature at the time (1–3). α-uL3 was raised against "cyL4" (Fleming et al., 1987). Our identifications of cyL4 and S-21 (original nomenclature) as α-uL3 and PSRP4/bTHXc (modern nomenclature), respectively, are based on correspondences of electrophoretic mobility in SDS-PAGE and pI values. In addition, PSRP4/bTHXc/S-21 was shown to be encoded by the nuclear genome, not the plastid genome (Fleming et al., 1987; Randolph-Anderson et al., 1989; Yamaguchi et al., 2002; Willmund et al., 2022). However, we retained the name "S-21" due to low uncertainty in this assignment. α-TOC159 antibody detects this protein because a Blastp search with the sequence of pea TOC159 (against which the antibody used here was raised) identified homologues in Chlamydomonas (e.g. Cre17.g734300; Score, 68.9; E value, 3e-11). TOC159 was detected as a 52 kDa truncated protein species which is generated by proteolytic cleavage of the full-length protein and remains membrane-inserted (Bauer et al., 2002; Joo et al., 2022).

**Supplemental Methods**

**Mass Spectrometry and proteomics.**

Proteins were concentrated in a stacking gel using SDS-PAGE stained with Coomassie Brilliant Blue R-250 (Biorad). Proteins were in-gel digested as follows. Proteins were reduced with 10 mM dithiothreitol (Sigma) in 50 mM $NH_4HCO_3$ (Sigma) for 30 min at RT and then alkylated with 50 mM iodoacetamide (Sigma) in 50 mM $NH_4HCO_3$ for 30 min at RT in dark. Gel pieces were washed with acetonitrile (ACN, BDH) and then rehydrated in trypsin digestion solution containing 25 mM

$NH_4HCO_3$ and 10 ng/µL of trypsin (Sigma) followed by incubation overnight at 30ºC. Tryptic peptides were extracted three times for 15 min at RT with extraction solution (60% acetonitrile + 0.5% formic acid (FA, Fisher), four volumes of the digestion solution). Peptides were dried using a Speedvac at 43 ºC and stored at -20 ºC until MS analysis. Liquid chromatography-tandem MS (LC-MS/MS) analyses were performed on a Thermo EASY nLC II LC system coupled to a Thermo LTQ Orbitrap Velos mass spectrometer equipped with a nanospray ion source. Tryptic peptides were resuspended in solubilization solution containing 97% of water, 2% of ACN and 1% of FA to give a peptide concentration of 100ng/µL. An aliquote (2 µL) of each sample was injected into a 10 cm × 100 µm column, which was in-house packed with Michrom Magic C18 stationary phase (5 µm particle diameter and 300 Å pore size). Peptides were eluted using a 90-min gradient at a flow rate of 400 nL/min with mobile phase A (96.9% water, 3% ACN and 0.1% FA) and B (97% ACN, 2.9% water and 0.1% FA). The gradient started at 2% of B, linear gradients of B were achieved to 8% at 16 min, 16% at 53 min, 24% at 69 min, 32% at 74 min, 54% at 81 min, 87% at 84 min followed by an isocratic step at 87% for 3 min and at 2% for 3 min. A full MS spectrum (m/z 400-1400) was acquired in the Orbitrap at a resolution of 60,000, then the ten most abundant multiple charged ions were selected for MS/MS sequencing in linear trap with the option of dynamic exclusion. Peptide fragmentation was performed using collision induced dissociation at normalized collision energy of 35% with activation time of 10 ms. Spectra were internally calibrated using polycyclodimethylsiloxane (*m/z* 445.12003 Da) as a lock mass.

Processing of MS data. MS data were processed using Thermo Proteome Discoverer software (v2.2) with the SEQUEST search engine. The database search was against the UniProt *Chlamydomonas reinhardtii* database (TaxID=3055, V2017-10-25). The enzyme for database search was chosen as trypsin (full) and maximum missed cleavage sites were set at 2. Mass tolerances of the precursor ion and fragment ion were set at 10 ppm and 0.6 Da, respectively. In the processing of the data of the proteomic analyses, static modification on cysteine (carbamidomethyl, +57.021 Da) and dynamic modifications on methionine (oxidation, +15.995 Da) and N-terminus (acetyl, +42.011 Da) were allowed. Proteins were identified with high confidence (false discovery rate <1%). For comparisons of LDM and TM by differential quantitative MS using metabolic labeling, the culture conditions are described above. In MS data processing, we allowed dynamic modifications on methionine (oxidation, +15.995 Da), on cysteine (carbamidomethyl, +57.021 Da), on protein N-termini (acetyl, +42.011 Da). For quantification of isotopically labeled peptides, following dynamic modifications were also included, on alanine, cysteine, aspartic acid, glutamic acid, phenylalanine, glycine, isoleucine, leucine, methionine, proline, serine, threonine, valine, tyrosine (1x label with $^{15}N$, +0.997 Da), on lysine, asparagine, glutamine, tryptophan (2x labels with 15N, +1.994 Da), on histidine (3x labels with $^{15}N$, +2.991 Da), and on arginine (4x labels with $^{15}N$, +3.988 Da). Relative protein abundances of isotopically labeled ($^{15}N$) and unlabeled control ($^{14}N$) samples were calculated with the precursor ion extracted-ion chromatogram areas using Thermo Proteome Discoverer software. Exp. q-values (combined) were less than 0.001. In both data sets, proteins were categorized manually based on annotations and the literature.

1. Schmidt, R. J., Myers, A. M., Gillham, N. W., and Boynton, J. E. (1984) Immunological similarities between specific chloroplast ribosomal proteins from Chlamydomonas reinhardtii and ribosomal proteins from Escherichia coli. *Mol Biol Evol*. **1**, 317–34
2. Randolph-Anderson, B. L., Gillham, N. W., and Boynton, J. E. (1989) Electrophoretic and immunological comparisons of chloroplast and prokaryotic ribosomal proteins reveal that certain families of large subunit proteins are evolutionarily conserved. *J Mol Evol*. **29**, 68–88
3. Banerjee, A. K., M, S., M, N., and Murty, U. S. (2010) Classification and clustering analysis of pyruvate dehydrogenase enzyme based on their physicochemical properties. *Bioinformation*. **4**, 456–62
